# Visualization of a multi-turnover Cas9 after product release

**DOI:** 10.1101/2024.11.25.625307

**Authors:** Kaitlyn A. Kiernan, David W. Taylor

**Affiliations:** Department of Molecular Biosciences, University of Texas at Austin, Austin, TX, USA; Institute for Cellular and Molecular Biology, University of Texas at Austin, Austin, TX, USA; Center for Systems and Synthetic Biology, University of Texas at Austin, Austin, TX, USA; Livestrong Cancer Institutes, Dell Medical School, Austin, TX, USA

## Abstract

While the most widely used CRISPR-Cas enzyme is the *S. pyogenes* Cas9 endonuclease (Cas9), it exhibits single-turnover enzyme kinetics which leads to long residence times on product DNA. This blocks access to DNA repair machinery and acts as a major bottleneck during CRISPR-Cas9 gene editing. Although Cas9 can eventually be forcibly removed by extrinsic factors (translocating polymerases, helicases, chromatin modifying complexes, etc), the mechanisms contributing to Cas9 dissociation following cleavage remain poorly understood. Interestingly, it’s been shown that Cas9 can be more easily dislodged when complexes collide with the PAM-distal region of the Cas9 complex or when the strength of Cas9 interactions in this region are weakened. Here, we employ truncated guide RNAs as a strategy to weaken PAM-distal nucleic acid interactions and still support Cas9 activity. We find that guide truncation promotes much faster Cas9 turnover and used this to capture previously uncharacterized Cas9 reaction states. Kinetics-guided cryo-EM enabled us to enrich for rare, transient states that are often difficult to capture in standard workflows. From a single dataset, we examine the entire conformational landscape of a multi-turnover Cas9, including the first detailed snapshots of Cas9 dissociating from product DNA. We discovered that while the PAM-distal product dissociates from Cas9 following cleavage, tight binding of the PAM-proximal product directly inhibits re-binding of new targets. Our work provides direct evidence as to why Cas9 acts as a single-turnover enzyme and will guide future Cas9 engineering efforts.

## Introduction

The CRISPR-Cas9 system (Clustered Regularly Interspaced Short Palindromic Repeats-CRISPR associated) from *Streptococcus pyogenes* carries out RNA-guided DNA target recognition and degradation and has been repurposed for precise genome manipulation^1,2^. Programmed with a single-guide RNA (sgRNA), Cas9 specifically targets and cleaves double-stranded DNA sequences flanked by a protospacer-adjacent motif (PAM). Following DNA cleavage, Cas9 remains stably associated with the dsDNA, limiting product release, and results in single-turnover cleavage kinetics (Figure 1a)^3,4^. This persistent product-bound state precludes access to the double-strand break (DSB) *in vivo,* preventing cellular DNA repair machinery, leads to slow DNA repair, and ultimately dictates repair outcomes^3,5–7^. While stable association can be beneficial for technologies that require prolonged Cas9 residence times, such as CRISPRi^8–13^, this slow dissociation acts as a barrier to achieving efficient gene editing *in vivo*.

**Figure 1.**
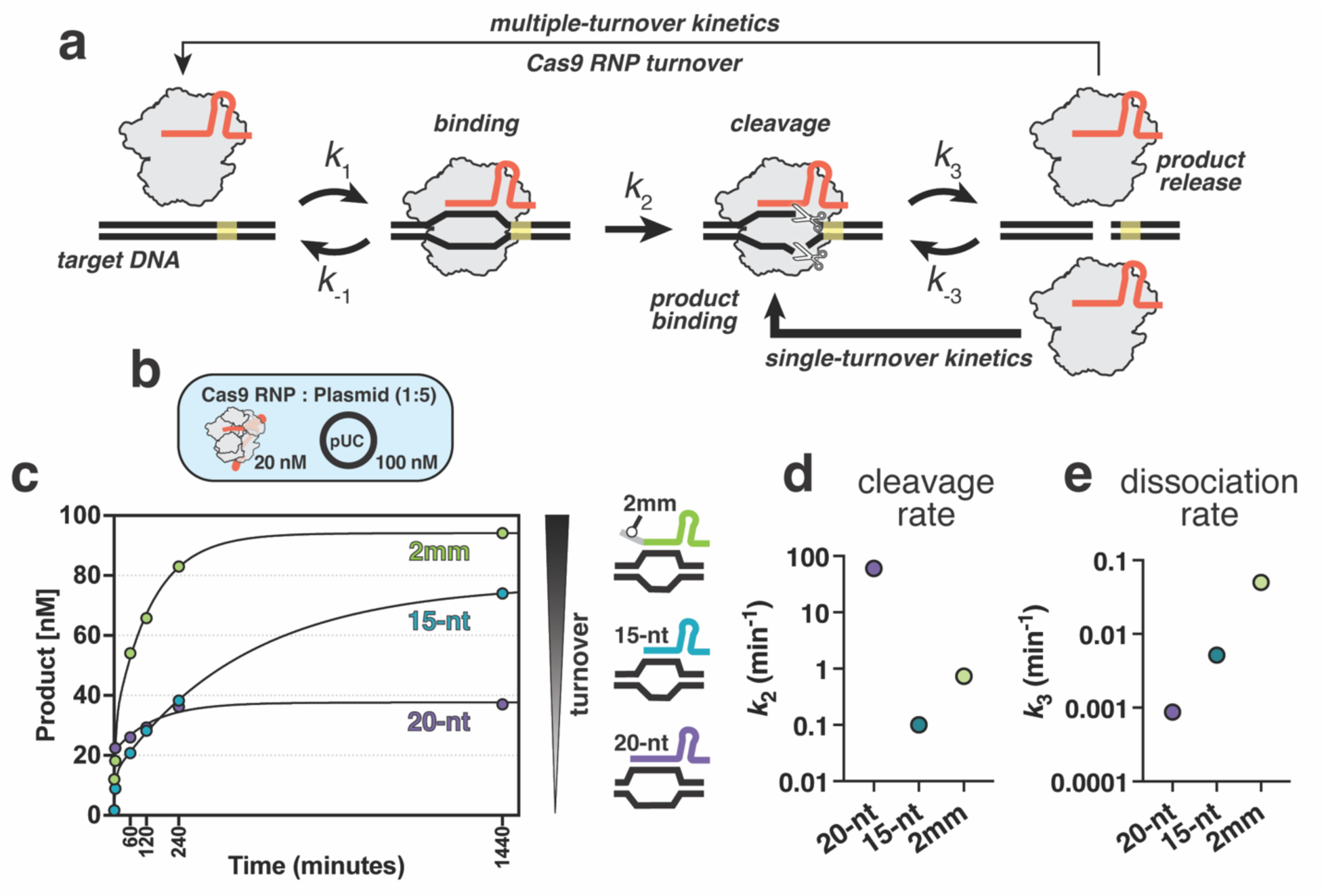
Kinetic analysis of Cas9 programmed with truncated sgRNAs. **(a)** Schematic of Cas9 reaction cycle. **(b)** Diagram of the experimental setup for Cas9 turnover assays where plasmid DNA was added to a pre-formed Cas9 RNP in 5-fold molar excess. **(c)** Cas9 turnover assays with 20-nt, 15-nt, and 2mm sgRNAs. 20 nM active Cas9 RNP was added to 100 nM plasmid substrate and product generation was plotted over time in minutes. **(d)** Cleavage rate constants (*k*_2_) derived from global fitting of the data shown in panel c. The cleavage rate constant for the 20-nt sample was locked at 60 min^-1^ as it was not captured in the time-scale used for the turnover assays and has been well-characterized in previous studies. **(e)** Dissociation rate constants (*k*_3_) determined for the 20-nt, 15-nt, and 2mm sgRNA samples.

Cas9 is a highly modular enzyme that undergoes large structural transitions during each stage of its reaction pathway, enabling conformational control of Cas9 activity^1,3,14–21^. Cas9 first binds to dsDNA targets via weak PAM interactions and triggers initial melting through structural distortion of the DNA. This melting enables Cas9 to probe for complementarity between the target strand (TS) of the DNA and spacer region of the sgRNA, proceeding in a directional manner towards the PAM-distal end. During R-loop propagation, the REC2 and REC3 domains are displaced from the central binding channel to accommodate the sgRNA-TS duplex, and once the R-loop proceeds past 14-bp, REC3 docks onto the PAM-distal region of the duplex^14,16,18^. Full accommodation of the R-loop in the central channel creates a kink in the DNA and ‘unlocks’ the HNH domain, enabling a 34 Å translation and ∼180° rotation towards the scissile phosphate at position 3 of the TS^19–21^. DNA unwinding and subsequent R-loop formation is the major energetic and kinetic barrier for Cas9 activation, and once this is complete, HNH and RuvC rapidly cleave the TS and NTS in a sequential, concerted manner^4,21–23^.

Cas9 activity being coupled to stable R-loop formation imparts an immense advantage for programmable gene editing and enables Cas9 activation when loaded with any sgRNA containing sufficient complementary to a PAM-containing DNA target. However, this stable R-loop formation prevents DNA re-winding following cleavage and leads to extremely slow dissociation rates, preventing release of DSBs and affecting Cas9 efficiency. Long Cas9 residence times have also been implicated in biasing DNA repair outcomes and is an important consideration for *in vivo* applications^6,7^. Several studies have shown that Cas9 dissociation requires translocating enzyme complexes, such as RNA polymerase, moving replication forks, and chromatin remodelers, to effectively dislodge Cas9 from a DSB^7,24,25^. It has also been shown that Cas9 dissociation occurs more readily if the collision with a translocating enzyme occurs on the PAM-distal side of Cas9^24,26,27^, however, this is no unifying mechanism describing Cas9 dissociation to support this.

To address this gap in the field, we use a combination of kinetics and cryo-electron microscopy (cryo-EM) to characterize the mechanisms controlling Cas9 turnover. By using kinetic analysis of Cas9 with altered guide-target duplexes, we identified a sgRNA truncation that resulted in increased Cas9 dissociation and multi-turnover cleavage kinetics. Cryo-EM analysis of a multi-turnover complex enabled us to capture snapshots of Cas9 throughout the entire reaction cycle, including the first direct visualization of Cas9 dissociating from the DNA product. We directly demonstrate that Cas9 turnover is limited by retention of the PAM-containing DNA product and occludes binding of new targets., Together, this work answers a long-standing question regarding exceptionally slow Cas9 turnover and offers a structural basis for Cas9 single-turnover kinetics.

## Results

### sgRNA truncation promotes faster Cas9 turnover

Cas9 turnover requires release of the heteroduplex, re-winding of the PAM-distal DNA and subsequent R-loop collapse. However, base-pairing between the sgRNA and DNA target promotes stable association of Cas9 and is required for conformational activation of the enzyme. Following DNA cleavage, Cas9 remains stably associated with the heteroduplex, preventing spontaneous Cas9 dissociation from DSBs. Eventually, Cas9 can be displaced by molecular motors or other enzymes that collide with the PAM-distal side of the Cas9-bound DSB^24,25,28^. Interestingly, mutations in high-fidelity Cas9 variants that remove REC3 contacts with the PAM-distal end result in higher Cas9 turnover with on-target DNA via increased DNA re-winding but exhibit lower overall cleavage rates than WT Cas9^29,30^. We reasoned that shortening the RNA:DNA hybrid would result in faster DNA rehybridization, R-loop collapse and subsequent turnover.

We and other groups have shown that sgRNA truncation down to a 14-nt spacer can support cleavage, albeit at much slower rates^31–34^. To test whether truncated sgRNAs promote multiple-turnover kinetics, we first benchmarked Cas9 cleavage efficiency of a fluorescently labeled 55-bp target DNA duplex when used with a 15-, 17-, or 20-nt spacer. When compared to the 17-nt and 20-nt sgRNAs, we observed that cleavage with the 15-nt sgRNA was greatly inhibited (Extended Data Figure 1). Even after a 20 hr incubation, we observed only a small fraction of the DNA substrate was cleaved.

DNA substrate topology and PAM-distal unwinding has been shown to modulate both cleavage efficiency with truncated sgRNAs and Cas9 displacement following DNA cleavage^35–37^. We hypothesized that these truncated sgRNAs would perform better with plasmid DNA and may be a better substrate for testing turnover activity. We then performed kinetic measurements of Cas9 cleavage of a target plasmid in 5-fold molar excess of Cas9. The full-length 20-nt sgRNA supported rapid cleavage of plasmid DNA but quickly exhibited a plateau of linear product corresponding to a stoichiometric amount of Cas9, confirming single-turnover kinetics behavior as previously reported (Figure 1c)^3,29,38^. In contrast, cleavage with the 15-nt sgRNA was much slower but continued without exhibiting a clear plateau and exhibited ∼2-fold increase in turnover (Figure 1c). This finding suggests that a shorter R-loop will be more likely to collapse following cleavage but is limited by the efficiency of catalytic activation.

We then explored how different guide-target combinations may retain turnover activity but increase cleavage rate. We hypothesized that destabilizing the 5’ end of the truncated sgRNA:DNA heteroduplex could increase the rate of DNA rehybridization in the PAM-distal region and, in turn, enhance turnover. To this end, we added one (1mm) or two (2mm) terminal mismatched nucleotides to the 5’ end of the 15-nt sgRNA and tested turnover activity. When compared to the fully hybridized 15-nt sgRNA, we observed enhanced turnover using both the 1mm and 2mm sgRNAs, with the highest being the 2mm sgRNA (Extended Data Figure 1). Excitingly, we found that the 2mm sgRNA was able to support near-complete cleavage of the 5-fold molar excess of plasmid substrate as well as rescue the cleavage defect observed for short, synthetic substrates as observed with the 15-nt sgRNA (Figure 1c, Extended Data Figure 1).

### Product inhibition limits Cas9 turnover

The Cas9 reaction pathway is comprised of multiple intrinsic rate constants which can be challenging to define from observed rates^39,40^. Here, we use global fitting to define the cleavage and dissociation rate constants from our kinetic measurements by fitting the data to a model that accounts for all the data (Figure 1a). For the 20-nt sgRNA, we observed the rate of product release (*k*_3_) was extremely slow (0.000871 min^-1^) and consistent with single-turnover kinetics. For the 15-nt sgRNA, we observed the slowest cleavage rate of 0.105 min^-1^ but with a dissociation rate (0.00517 min^-1^) approximately 10-fold faster than product release with a full-length guide (Figure 1c). The 2mm sgRNA supported rapid product release with a dissociation rate of 0.0501 min^-1^, 60-fold faster than a 20-nt sgRNA, compensating for the reduced cleavage rate of 0.731 min^-1^ (Figure 1c).

Altogether, our kinetic analysis suggests that sgRNA truncation does promote product release following cleavage and that mismatches in the 5’ region of the sgRNA:DNA hybrid favor rehybridization of the TS with the NTS post-catalysis, inducing spontaneous Cas9 dissociation from the product. With 20-nt sgRNAs, extremely slow product release limits Cas9 turnover, and as the reaction equilibrium shifts towards a higher concentration of the product relative to the substrate, is more likely to be bound to product than substrate, providing a potential molecular mechanism for long Cas9 residence times on a DSBs.

### Kinetics-guided cryo-EM provides snapshots spanning the entire reaction cycle

We next sought to determine the molecular basis for Cas9 dissociation from DNA using cryo-electron microscopy (cryo-EM). Heterogeneity in cryo-EM datasets can be problematic when averaging particles to generate enough signal to produce a high-quality reconstruction because while averaging offers a net gain in resolution, high degrees of motion and heterogeneity leads to loss of information in regions that are often the most interesting. Accordingly, samples are typically prepared to maximize homogeneity to determine a single, high-resolution structure.

Cas9 is particularly susceptible to the heterogeneity problem as many domains undergo large structural changes during the reaction and are typically poorly resolved. To circumvent this, many studies have used catalytically inactive complexes or modified DNA substrates to trap Cas9 in specific conformations^18,21,41,42^ but as these represent off-pathway intermediates, we decided to implement a different approach. This strategy involves using kinetics to guide cryo-EM sample preparation and has been previously used by our group to yield datasets capturing multiple on-pathway intermediate Cas9 structures under conditions that support catalytic activity^20,43^. By carefully selecting timepoints for vitrification, we enrich for maximal heterogeneity to capture rare, transient intermediates that are otherwise missed. As Cas9 dissociation is slow, this is an extremely rare state and structural details regarding dissociation following DNA cleavage remain uncharacterized. Faster dissociation kinetics observed with our mm2 sgRNA sample made it the perfect candidate to visualize Cas9 in the process of dissociating from its product.

Based on our kinetic analysis with the mm2 complex, we hypothesized that a timepoint mid-way through the reaction would capture Cas9 during all stages of the reaction and chose to prepare grids with samples incubated for 2 hours at 37°C. To classify the conformational states existing in our dataset, we used multiple rounds of 3D classification implemented in cryoSPARC^44^ and generated an ensemble of reconstructions comprising a total of 25 subclasses with resolutions ranging from 2.8 - 3.4 Å. (Figure 2b). Each subclass was manually inspected and sorted into five distinct conformational states (Figure 2b, Extended Data Figure 4). These states represent the progression of Cas9 through its reaction cycle and comprise the (I) pre-activation (II) pre-cleavage (III) checkpoint (IV) product and (V) dissociated states (Figure 2c).

**Figure 2.**
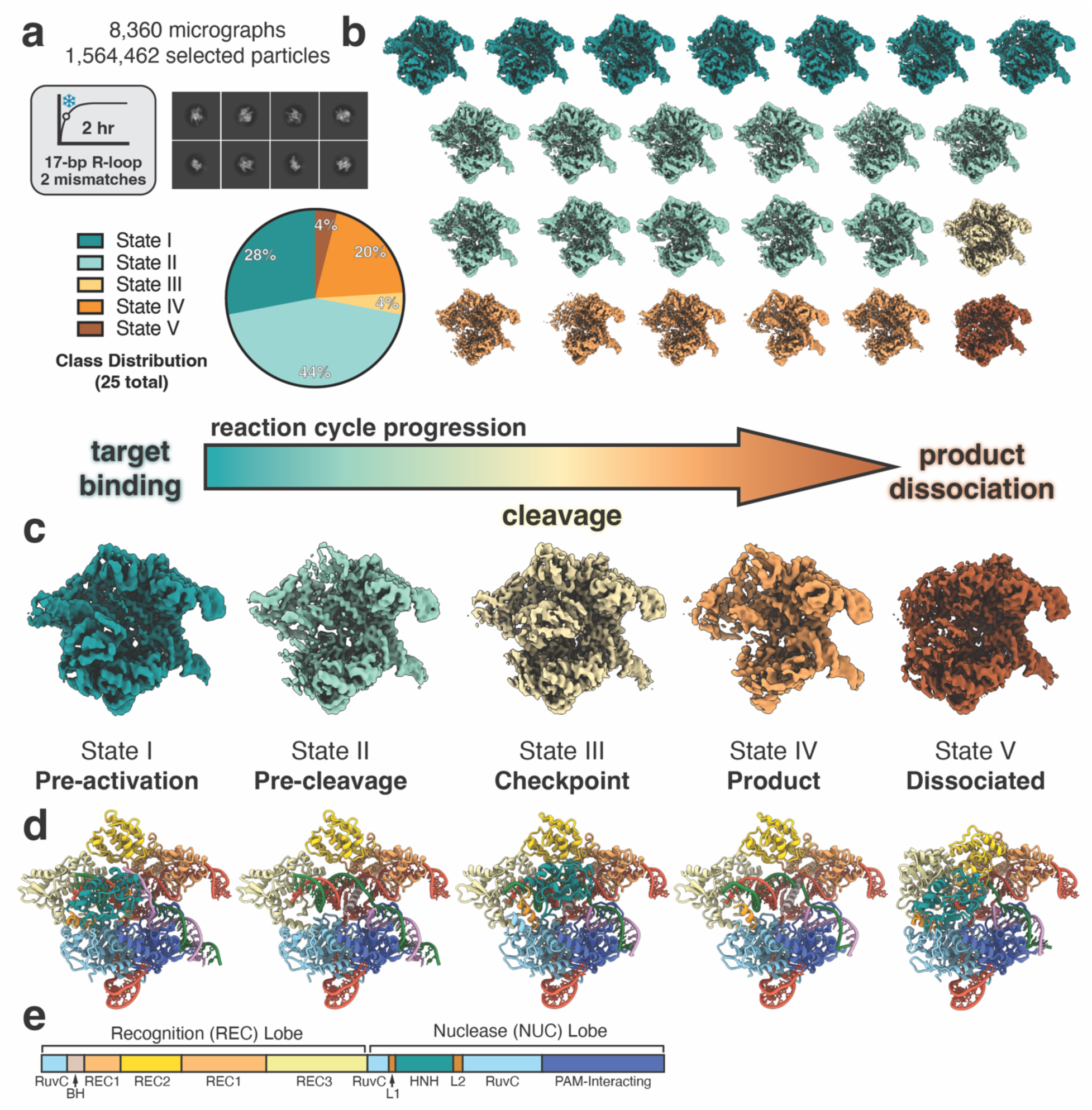
Kinetics-guided cryo-EM data analysis. **(a)** Data collection conditions used for cryo-EM. The pre-formed Cas9 RNP containing a 17-bp R-loop with 2 terminal mismatches was incubated with a 55-bp dsDNA target for 2 hours at 37C before vitrification. Representative 2D classes are shown from the final particle stack used for further 3D classification. 3D class distribution represented as a pie chart. **(b)** Ensemble of cryo-EM reconstructions generated using 3D classification. **(c)** Representative maps corresponding to five main conformational states that span the entire reaction cycle from target binding to product dissociation. **(d)** Structures of each state corresponding to the maps shown in panel c. Colored according to the schematic below. **(e)** Cas9 architecture colored by domain.

In the pre-activation state, Cas9 is bound to the target DNA where the R-loop has been fully formed but the PAM-distal DNA has not been fully accommodated into the central channel (Extended Data Figure 3). In this state, HNH remains docked onto the RuvC domain and the DNA is in a linear conformation (Figure 3a,d). Following R-loop completion, the DNA adopts a kinked conformation and induces large conformational changes in the nuclease domains. The pre-cleavage state represents this intermediate where we observe that the DNA is bent towards the central channel, diffuse density for both HNH and RuvC nuclease domains indicates rearrangement towards active conformations, and the DNA remains intact (Figure 3a,d).

**Figure 3.**
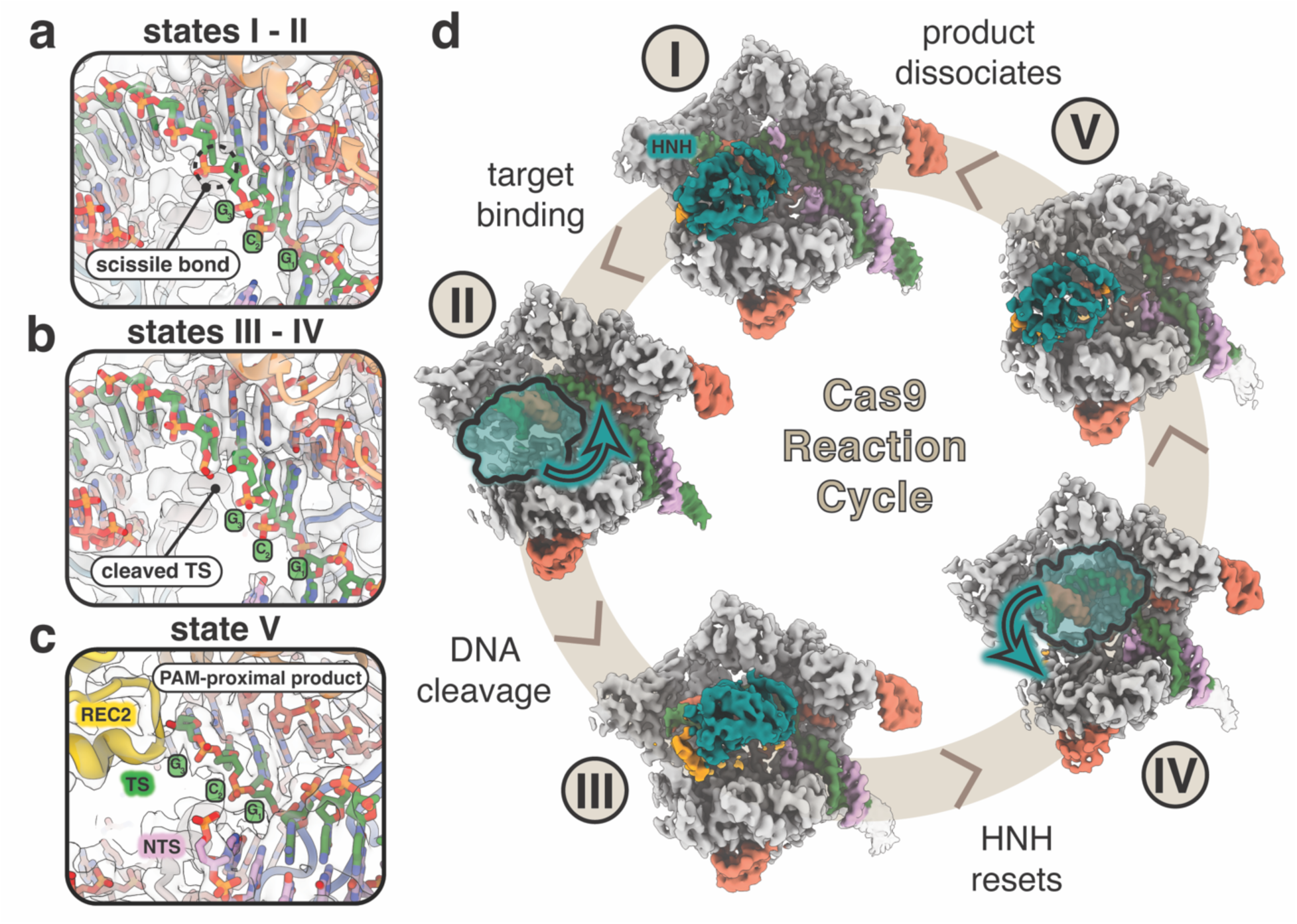
Structures of Cas9 spanning the entire reaction from a single dataset. **(a)** Close-up view of the intact scissile bond of the target strand (TS) in states I and II. Map density shown from state II. **(b)** Close-up view of the cleaved scissile bond showing clear separation between nucleotides 3 and 4 of the hybridized TS. Map density shown from state III. **(c)** Close-up view of the cut site in state V following PAM-distal product release. PAM-proximal product remains bound where the first three nucleotides of the TS upstream of the PAM remain hybridized. **(d)** The five main conformational states populate the entire reaction cycle where (I) represents the pre-activation state just following R-loop completion, (II) represents the pre-cleavage state during reorganization of the NUC lobe on-path to dock at the cleavage sites, (III) represents the conformational checkpoint state where HNH undocks following DNA cleavage, (IV) represents the product state where HNH remains flexibly tethered after undocking from the cut TS, and (V) represents the dissociated state where the PAM-distal end of the DNA is released from the complex.

### The HNH domain resets following DNA cleavage

State III closely resembles the conformational checkpoint state (PDB 7Z4L, RMSD 0.796 Å) where the HNH domain has repositioned onto the TS:sgRNA heteroduplex but the scissile bond is still ∼30 Å away from the active site (Extended Data Figure 2). However, our structure has a critical difference where we observe the TS has been cleaved, evidenced by unambiguous density in our high-resolution map (Figure 3b). This provides direct evidence that following DNA cleavage, the HNH domain dissociates from the target strand (TS) and resets to adopt the checkpoint conformation once again (Extended Data Figure 2). In our product state structure, we see that the nuclease domains become largely disordered following release of the heteroduplex, indicating that this is an intermediate following DNA cleavage where HNH remains highly flexible (Figure 3b). A previous single-molecule FRET (smFRET) study has shown that following DNA cleavage, HNH adopts the conformational checkpoint state again^45^, but was later confounded by a separate smFRET study showing HNH is highly flexible following cleavage^46^. Our findings reconcile both FRET studies by directly showing that after TS cleavage, the HNH domain does reset to the conformational checkpoint state and subsequently displays a high degree of flexibility. Given the timescale of these FRET studies (20 s) is much shorter than the lifetime of a Cas9 product complex (minutes to hours), our structural data provides further insight into longer-term dynamics of the HNH conformational landscape following DNA cleavage.

REC3 normally docks onto the 15-20 bp region of the distal duplex and imparts a kink in the DNA, facilitating the reorganization of the NUC lobe. In other product state structures, REC3 remains docked in this region^19–21^. In contrast, we find that in our product state structure, the REC3 domain has undocked from the PAM-distal duplex, moving away from the central binding channel and releasing the DNA kink to form a linear duplex once again (Extended Data Figure 3). The PAM-distal duplex is also stabilized by electrostatic interactions within the RuvC domain, including a flexible, positively-charged loop that is normally only resolved in product state structures (Extended Data Figure 3)^19^. In our product state, these residues are disordered and are not observed to establish the same contacts with the distal DNA backbone. Without stabilization by REC3 and RuvC contacts, the PAM-distal duplex exhibits a higher degree of flexibility following DNA cleavage and likely contributes to faster DNA rewinding and R-loop collapse.

### Cas9 dissociates from the PAM-distal DNA product

Remarkably, in the dissociated state (state V), we observe particles in which the Cas9 complex reverts almost entirely back to its binary conformation (Figure 4e). Excitingly, the PAM-distal DNA product has been released from the complex. The REC2 and REC3 lobes move closer to the center of the complex and once again occlude the central DNA binding channel (Figure 4e). While the sgRNA seed (1-10 nt) remains preordered for binding, the PAM-containing half of the product remains stably bound (Figure 4b, d). Two residues, R1333 and R1335, in the PAM-interacting region directly participate in base-specific hydrogen bonding with the GG dinucleotides in the PAM and are maintained in the dissociated state. This persistent PAM-binding directly interferes with binding new targets and explains why Cas9 exhibits single-turnover kinetics.

**Figure 4.**
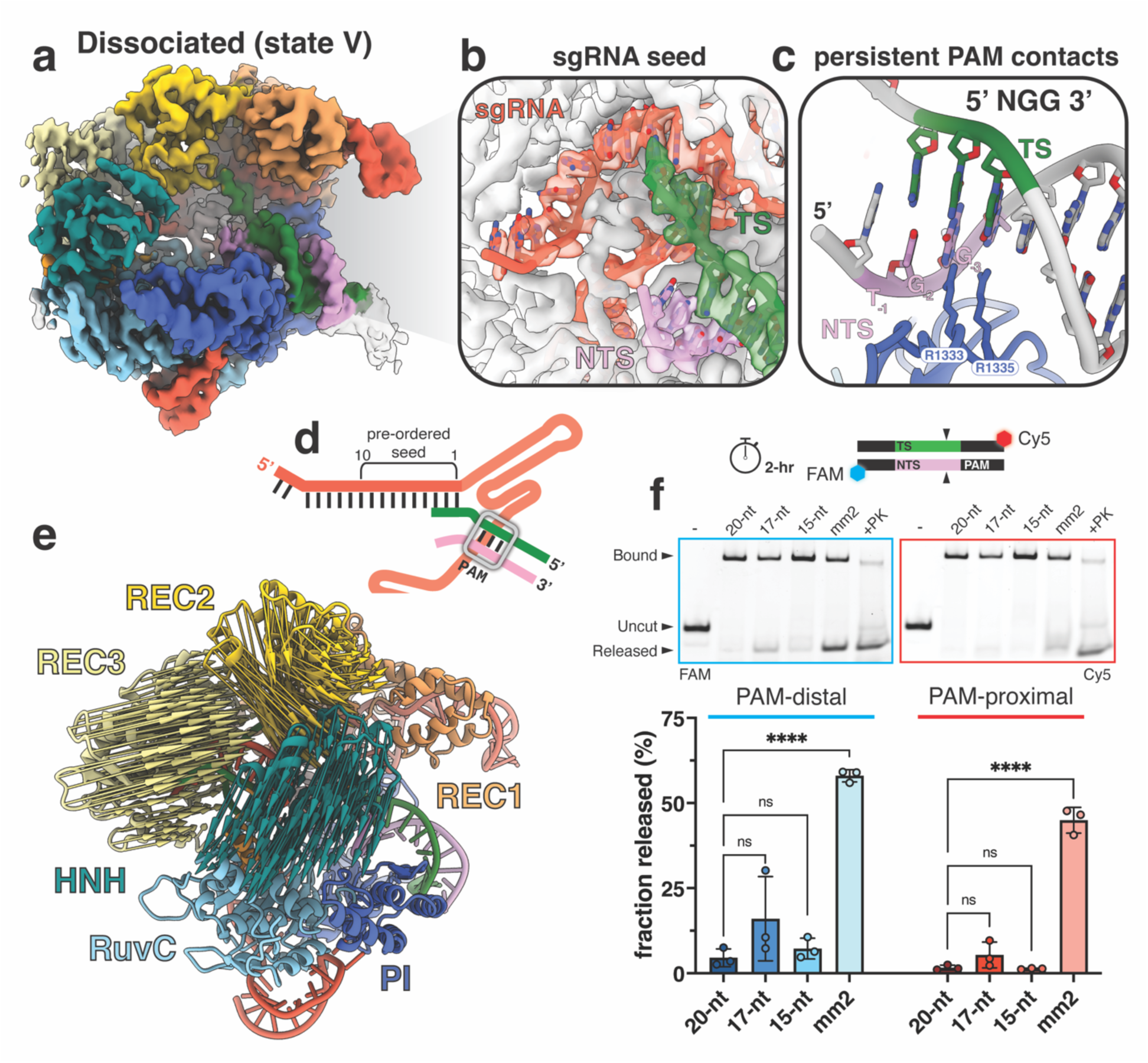
Structural basis for slow Cas9 dissociation. **(a)** Cryo-EM map of the dissociated state colored by domain. **(b)** Zoom-in of the pre-ordered sgRNA seed region. **(c)** Zoom-in view of the PAM contacts maintained even following PAM-distal product release. **(d)** Schematic of the nucleic acid in this complex. sgRNA colored in red, target strand in green, and non-target strand in pink. PAM site indicated with gray box. **(e)** Modevector arrows showing the domain movements from the checkpoint to dissociated states. **(f)** Monitoring PAM-distal and PAM-proximal product release following cleavage. Cas9 programmed with a 20-, 17-, 15-nt, or 2mm sgRNA was incubated with target DNA labeled where the PAM-distal end was labeled with FAM and the PAM-proximal end labeled with Cy5. Product release was analyzed at 2 hours following reaction initiation on Native 4-20% polyacrylamide gels. Samples were quantified using ImageJ and plotted as a ratio of released product to bound product and represented as a percentage. Data represented as the mean ± s.d. from three independent replicates and statistical significance determined with using a one-way ANOVA, \*\*\*\**P<0.0001*.

Product dissociation and subsequent re-binding are difficult to deconvolute *in vitro*, especially when utilizing plasmid substrates. We hypothesized that this phenomenon could be more clearly observed with linear fragments given our observation of Cas9 dissociation in our structural dataset where we used short, linear substrates. To assess PAM-distal and PAM-proximal product dissociation *in vitro*, we labeled the ends of a target DNA with different fluorophores and monitored product release with different sgRNAs after 2 hours (Figure 4f). When programmed with a 20-nt or 17-nt sgRNA, consistent with single-turnover kinetics, Cas9 tightly retains both ends of the DNA product. Strikingly, we observe significantly more product release of both DNA ends when Cas9 is programmed with the 2mm sgRNA, exhibiting over a 50-fold increase in product release of the PAM-distal end of the duplex (Figure 4f).

Our observations align with previous studies showing that the PAM-distal non-target strand (NTS) is released and accessible after cleavage^25^, likely contributing to rehybridization with the PAM-distal TS and subsequent dissociation of the PAM-distal end of the DNA. Our findings that the PAM-proximal product remains tightly associated, particularly when canonical sgRNAs are used, provides the structural and kinetic basis for slow turnover. To our knowledge, this is the first direct visualization of Cas9 following product dissociation and contributes to an updated model for the Cas9 reaction cycle (Figure 5). Persistent binding of Cas9 to the PAM site emphasizes the importance of this interaction in the enzyme’s turnover dynamics and highlights a potential target for improving the efficiency of CRISPR-Cas9 gene editing.

**Figure 5.**
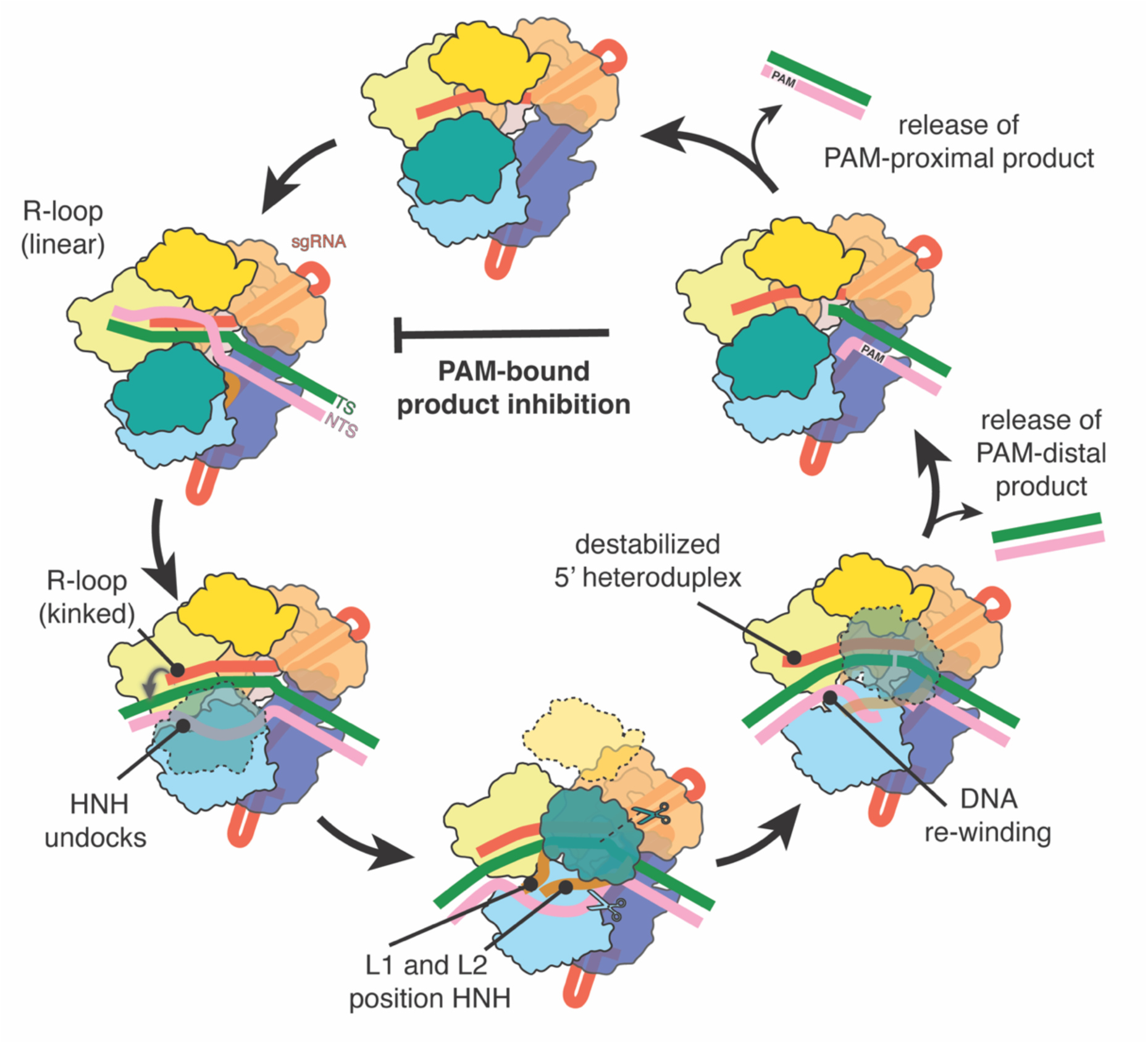
Updated model of Cas9 reaction cycle. The cycle begins with a Cas9-sgRNA binary complex searching for PAM-containing DNA targets. Complementarity of the sgRNA spacer and target sequence initiates R-loop propagation where the distal duplex adopts a linear conformation. When the R-loop is fully formed, the PAM-distal DNA duplex adopts a kinked conformation and initiates nuclease lobe reorganization. The L1 and L2 linker helices precisely position HNH onto the target strand. HNH reorganization exposes the RuvC active site and enables accommodation of the NTS. HNH and RuvC catalyze concerted TS and NTS cleavage, respectively. Following TS cleavage, the HNH domain moves in reverse and adopts the checkpoint conformation again. Coordinated cleavage of the NTS allows for PAM-distal DNA rewinding and triggers the slow release of the PAM-distal product. The PAM-proximal product remains bound to the complex, prevents re-binding of a new target, and leads to product inhibition. Eventual release of the PAM-proximal product resets Cas9 to the binary surveillance complex and restarts the cycle.

## Discussion

Our work answers a long-standing question in the field, explaining why Cas9 is a single-turnover enzyme. We demonstrate that persistent binding of the PAM-containing DNA product inhibits Cas9 turnover and prevents binding to new targets. We discovered that decreasing the sgRNA spacer length and PAM-distal heteroduplex stability can promote R-loop collapse following DNA cleavage and transforms Cas9 into a multi-turnover enzyme. We leveraged our kinetic information to capture 25 high-resolution snapshots of Cas9 in action, including the first visualization of Cas9 dissociated from product DNA. By defining the mechanisms for Cas9 product release and subsequent turnover, our work solidifies a comprehensive, unifying model for the Cas9 reaction cycle (Figure 5).

A multi-turnover Cas9 overcomes a significant bottleneck in the gene editing cycle and carries immense potential for biotechnology applications. Multi-turnover Cas9 complexes would no longer require cellular machinery to remove it from a DSB and subsequent repair would happen much faster. Persistent Cas9-DSB interactions can also modulate repair outcomes, skewing repair towards NHEJ^44^. HDR-mediated repair is limited by low efficiency and has been addressed by mutating or inhibiting genes involved in other DNA repair pathways^47,48^. It is plausible that a multi-turnover Cas9 would accelerate rates of HDR-mediated repair, circumventing the need to suppress other repair pathways and thus expanding upon available technologies for precision editing. Truncated sgRNAs have also been shown to increase editing fidelity *in vivo.* Our findings that a truncated sgRNA with two mismatches enhances turnover emphasizes the importance of assessing off-target editing when employing truncated sgRNAs *in vivo*.

Many Cas9 re-engineering strategies have focused on designing Cas9 variants with expanded PAM recognition to increase the number of targets that are accessible for editing^49–52^. One PAM-less variant, SpRY-Cas9 (SpRY), was engineered to harbor 11 mutations in the PAM-interacting domain and can recognize all potential PAM sites^52^. Additional non-specific electrostatic interactions increase binding affinity for DNA in a sequence-independent manner and enables SpRY to interrogate many more sites in the genome, but this results in slow target search and accumulation at off-target sites^43^. Our finding that Cas9 remains tightly associated with the PAM, even after partial DNA release, suggest that SpRY may not only exhibit slow target search but likely retains even tighter binding to the product following cleavage, offering an additional explanation for its reduced efficiency. A similar strategy was recently employed to engineer two naturally high-fidelity Cas9 enzymes, PsCas9 and FnCas9. Increasing Cas9 binding affinity to target DNA improved editing efficiency *in vivo,* but introduction of new electrostatic interactions with the PAM-proximal DNA may likewise impact turnover kinetics.

Overall, our study expands upon our current understanding of Cas9 biology and offers a novel strategy to turn Cas9 into a multi-turnover enzyme. We were able to interrogate the conformational landscape of Cas9 throughout the entire reaction cycle and uncover the elusive molecular mechanisms contributing to slow turnover of Cas9. This work provides critical insight that will guide both careful selection of the appropriate Cas9-based tool and the future design of new editors.

## Methods

### Cas9 expression and purification

*S. pyogenes* Cas9 (Cas9) constructs were cloned into a pET28b expression vector containing a C-terminal His_6_ tag. Recombinant Cas9 was expressed in *Escherichia coli* strain OverExpress C41(DE3) (Sigma) and purified by Ni-NTA (nickel-nitrilotriacetic acid) and ion exchange chromatography (IEX). Expression was induced when cells reached an OD_600_ of ∼0.6 by adding isopropyl β-D-1-thiogalactopyranoside (IPTG) to a final concentration of 0.2 mM. Cells were incubated for 18–20 hours at +18°C while shaking and then pelleted at +4°C by centrifugation at 7,000 rpm for 30 minutes. The cell pellet was resuspended in lysis buffer (20 mM HEPES-NaOH, pH 7.5, 300 mM NaCl, 3 mM βME, 10% (v/v) glycerol) supplemented with a protease inhibitor tablet (Roche). Cells were sonicated on ice and the lysate was clarified at 15,000 rpm for 45 minutes. Lysate supernatant was loaded onto a His-Trap FF crude column (Cytiva), washed with the lysis buffer containing 20 mM imidazole, and eluted with the same buffer but supplemented with 350 mM imidazole. Subsequent purification was performed using a HiTrap SP Sepharose High Performance cation exchange column (Cytiva) and a linear gradient of 100–500 mM KCl in 20 mM HEPES-KOH, pH 7.5. Fractions containing Cas9 were pooled, concentrated to 10 mg×mL^-1^, and buffer exchanged into Cas9 protein storage buffer (20 mM HEPES, pH 7.5, 200 mM NaCl, 1 mM EDTA, 10% (v/v) glycerol, 0.5 mM TCEP). All purified samples were aliquoted into single-use volumes, flash-frozen and stored at -80°C.

### Nucleic acid preparation

The 55-bp DNA substrates were prepared by annealing two complementary ssDNA oligonucleotides (IDT). Each oligo was resuspended in water to a final concentration of 100 μM. Both oligonucleotides were combined a 1:1 molar ratio and annealed by heating to +95°C for 5 minutes and slowly cooling to room temperature. All sgRNAs used in this study were purchased from GenScript, resuspended in nuclease-free water to a final concentration of 20 μM, flash-frozen in liquid nitrogen and stored at - 80°C. Sequences for oligonucleotides and sgRNAs listed in Extended Data Table 2. Fluorescently labeled substrates were prepared by annealing a 5’ 6-FAM TS oligo with an unlabeled NTS oligo. For the plasmid cleavage assays, the same Cas9 target site that was used in the linear fragments was cloned into pUC19, prepared in bulk via the ZymoPURE II Plasmid Maxiprep kit, and concentrated with the Vacufuge Plus Vacuum Concentrator (Eppendorf).

### Cryo-EM Sample Preparation and Data Collection

Cas9 complex samples were prepared in complex buffer (20 mM HEPES, pH 7.5, 200 mM NaCl, 1.0 mM EDTA, 10 mM MgCl_2_, 0.5 mM TCEP) in a 1:1.2:1.2 molar ratio of Cas9:sgRNA:DNA. WT SpCas9 and the sgRNA were combined and incubated for 10 minutes at room temperature before adding the 55-bp dsDNA substrate to initiate the reaction. The ternary complex was incubated for 2 hours at 37°C and the reaction was quenched via vitrification. 2.5 μL of sample was applied to glow-discharged holey carbon grids (Quantifoil 1.2/1.3), blotted for 7 s with a blot force of 0 and rapidly plunged into liquid nitrogen-cooled ethane using an FEI Vitrobot MarkIV. Data were collected on an FEI Titan Krios cryo-electron microscope equipped with a K3 Summit direct electron detector (Gatan). Images were recorded with SerialEM with a pixel size of 0.8332 Å at 13.3 electrons/pixel/second for 6 s (80 frames) to give a total dose of 80 electrons/pixel.

### Cryo-EM Data Processing

All data was processed in cryoSPARC v4.0^44^ and the overall workflow is included in Extended Data Figure 5. CTF correction, motion correction and template-based particle picking were performed in real-time using cryoSPARC Live v4.0.0. Particles were subject to multiple rounds of 2D classification and junk particles were filtered using multi-class ab-initio reconstruction and subsequent heterogenous refinement. The filtered particle stack was subject to multiple rounds of 3D classification (10 classes per round) using the PCA initialization mode with forced hard classification. One final round of non-uniform refinement^53^ was conducted using per-particle defocus and global CTF optimization parameters to generate the final 3D reconstructions. Cryo-EM map quality and model refinement statistics included in Extended Data Table 1 and Extended Data Figure 6.

### Atomic model building and refinement

To build the atomic models, structures 6O0Z, 7S4V, 7Z4L, 7S4V, and 4ZT0 were rigid body fit into the pre-activation, pre-cleavage, conformational checkpoint, product, and dissociated states, respectively. 3D models were manually adjusted and inspected in Coot ^54^ and ISOLDE ^55^, and structures were real-space refined in Phenix^56^. All figures were generated in ChimeraX v1.8^57^. Statistics for data collection and model refinement are reported in Extended Data Table 1. Modevector arrows between aligned models were generated using PDBarrows as described in Chaaban et al^58^.

#### Cas9 Cleavage Assays

Cleavage reactions were assembled in 1X cleavage buffer (20 mM HEPES, pH 7.5, 100 mM KCl, 0.5 mM TCEP, 10 mM MgCl2, 5% (v/v) glycerol). To form the Cas9-RNP, recombinant Cas9 was mixed with pre-annealed sgRNA in a 1:1.5 ratio and incubated at room temperature for 15 min. dsDNA was added to initiate the reaction. For turnover assays, DNA was added in a 5:1 molar ratio (dsDNA:Cas9-RNP). At each timepoint, an aliquot of the reaction was quenched by addition of 500 mM EDTA and 20 μg Proteinase K (Thermo Scientific). Cas9 cleavage products using fluorescently labeled 55-bp linear fragments were resolved on a 15% (w/v) TBE-Urea (7 M) polyacrylamide gel and ran at 180 V for 45 minutes. Plasmid cleavage products were resolved on a 1% agarose gel in 1X TAE buffer ran at 90 V for 75 min and post-stained with GelRed (Milipore Sigma). All gels were imaged using the ChemiDoc MP (Bio-Rad). Quantification of cleaved products was determined by densitometry analysis in ImageJ^59^. Linear plasmid product was normalized to EcoRI-digested plasmid DNA.

#### Kinetic Analysis

Kinetic measurements of Cas9 cleavage reactions were subject to global fitting in KinTek Explorer. Turnover experiments were fit to the reaction scheme in in Figure 1e where:

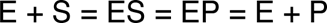

Experimental details of reactant concentrations were input for each experiment. In fitting data by simulation, each experiment is modeled exactly as it was performed. For the second-order DNA binding step (k_1_), the rate was not defined by the data and the binding rate constant and the reverse rate constant were locked at values to give a K_d_ for DNA binding of 1 nM, similar to other estimates for the equilibrium constant for Cas9 binding to DNA^20,4,30^. The cleavage rate constant for the 20-nt sample was also locked at 60 min^-1^ as it was not captured in the time-scale used for the turnover assays and has been well-characterized in previous studies^4,20,30^.

## Data and code availability

The atomic coordinates have been deposited in the Protein Data Bank (PDB) under the accession numbers 9EAK (pre-activation), 9EAL (pre-cleavage), 9ED9 (conformational checkpoint), 9EDA (product), and 9EDB (dissociated). Cryo-EM maps have been deposited in the Electron Microscopy Data Bank (EMDB) under accession numbers EMD-47834 (pre-activation), EMD-47835 (pre-cleavage), EMD-47941 (conformational checkpoint), EMD-47942 (product), and EMD-47943 (dissociated). Source data are available.

## Acknowledgements

We thank Dr. Kenneth Johnson for assistance with kinetic analysis and helpful discussion as well as Dr. Jack Bravo and members of the Taylor lab for insightful comments on the manuscript. Data were collected at the Sauer Structural Biology Laboratory at the University of Texas at Austin. This work was supported in part by the Welch Foundation grant F-1938 (to D.W.T.), the National Institutes of Health grant R35GM138348 (to D.W.T.), and a Robert J. Kleberg, Jr. and Helen C. Kleberg Foundation Medical Research Grant (to D.W.T.). The content is solely the responsibility of the authors and does not necessarily represent the official views of the National Institutes of Health.

## Notes

### Competing Interest Statement

The authors have declared no competing interest.

