## Extended Data for "Visualization of a multi-turnover Cas9 after product release"

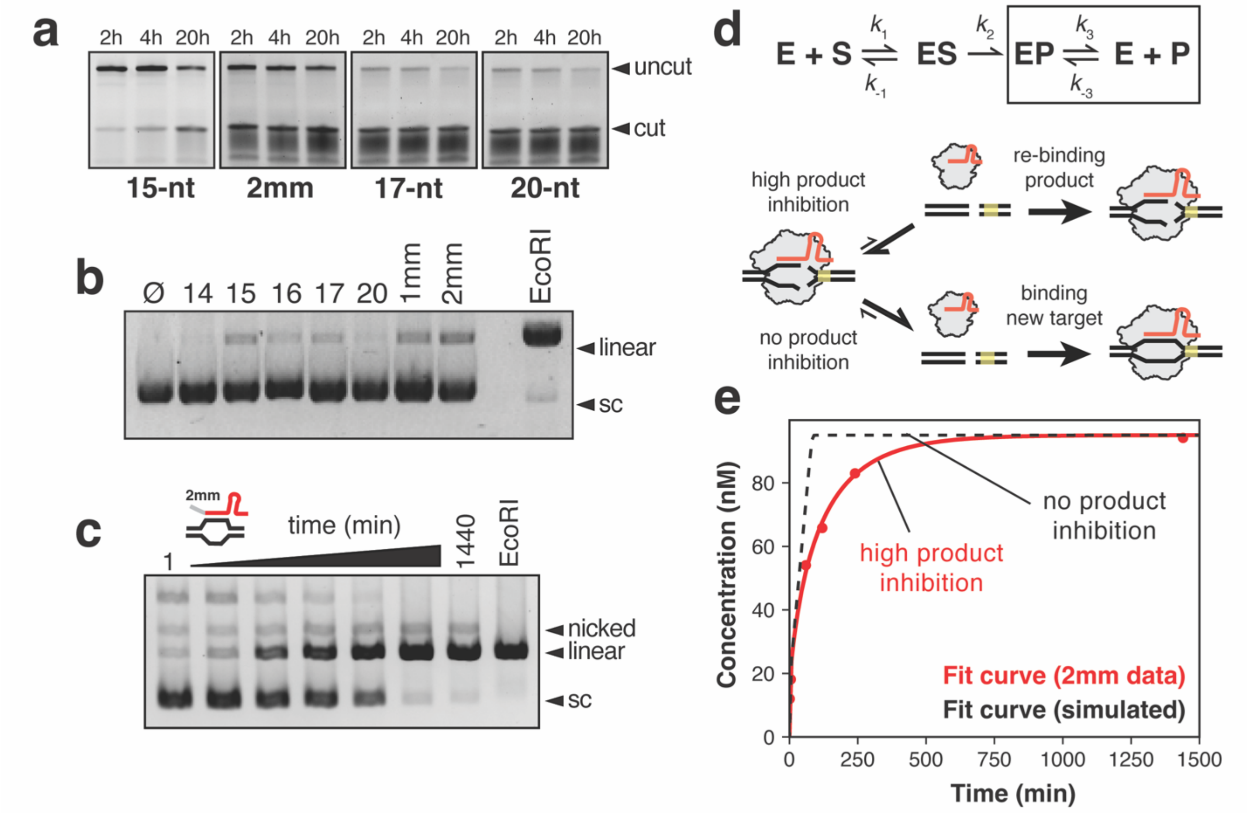


**Extended Data Figure 1. Kinetic analysis of Cas9 programmed with truncated sgRNAs. (a)** Cleavage of a fluorescently labeled 55-bp dsDNA target with a 15-nt, 2mm, 17-nt, and 20-nt sgRNA. Cleavage products were analyzed on 15% polyacrylamide denaturing gels. **(b)** Cas9 cleavage assay with plasmid DNA in high excess. Linear product measured following 18 hours of incubation with 14-, 15-, 16-, 17-, and 20-nt sgRNAs, and with a 16-nt sgRNA containing one mismatch (1mm) and 17-nt sgRNA with two terminal mismatches (2mm). **(c)** Turnover assay with the 2mm sgRNA sample. 20 nM active Cas9 RNP was added to 100 nM plasmid substrate and product generation was measured at 1, 5, 60, 120, 1440, and 2880 minutes. Fully-digested plasmid with EcoRI used as a control. Cleavage products were visualized using 1% agarose gels post-stained with 3X GelRed. **(d)** Equation used to fit kinetic data with box around the product release step of the reaction. Schematic below showing difference in reaction outcome with high degrees of product inhibition or no product inhibition present within the reaction. **(e)** Kinetic curves of experimental data (2mm) displaying high degrees of product inhibition overlayed with simulated data that have minimal product inhibition.

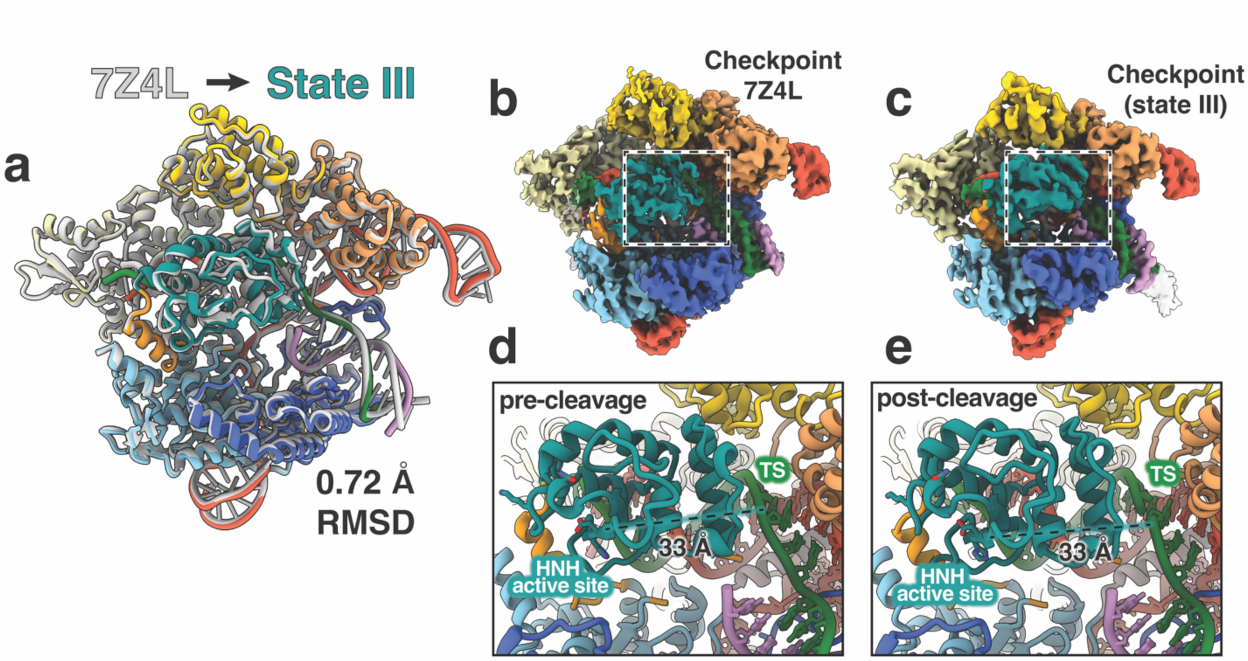


**Extended Data Figure 2. Analysis of the conformational checkpoint state. (a)** Overlay of PDB 7Z4L with the conformational checkpoint state (state III) in our dataset. **(b-c)** Cryo-EM maps colored by domain of the checkpoint state from **(b)** Pacesa et al 2022 and **(c)** this study. **(d)** Checkpoint structure before docking onto the TS for cleavage. The active site residues shown as sticks and is ~ 33 Å away from the TS. **(e)** Checkpoint structure directly following DNA cleavage after HNH is undocked from the TS. HNH adopts the same conformation as seen before cleavage with the active site rotated away ~ 33 Å from the cleaved TS.

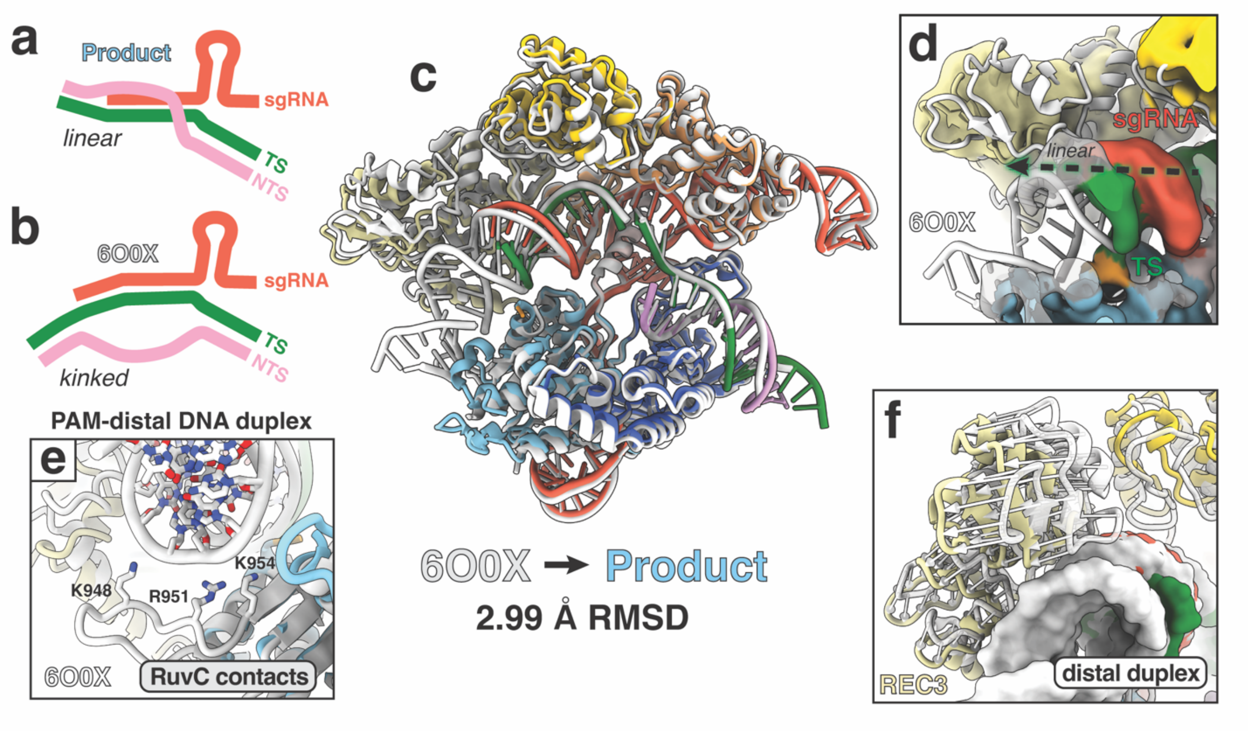


**Extended Data Figure 3. Structural comparison of product state. (a)** Schematic of the linear R-loop. **(b)** Schematic of the kinked R-loop. **(c)** Overlay of PDB 6O0X with the product state from our dataset. **(d)** Low-pass filtered product state map (6 Å) shows the path of the DNA duplex follows a linear trajectory compared with the kinked trajectory observed in 6O0X. **(e)** In 6O0X, electrostatic interactions in the RuvC domain are established with the PAM-distal DNA. These interactions are not observed in our product state structure. **(f)** Modevector arrows showing REC3 movement away from the heteroduplex in our product state (yellow) whereas in the 6O0X structure, REC3 remains docked onto the PAM-distal heteroduplex (white).

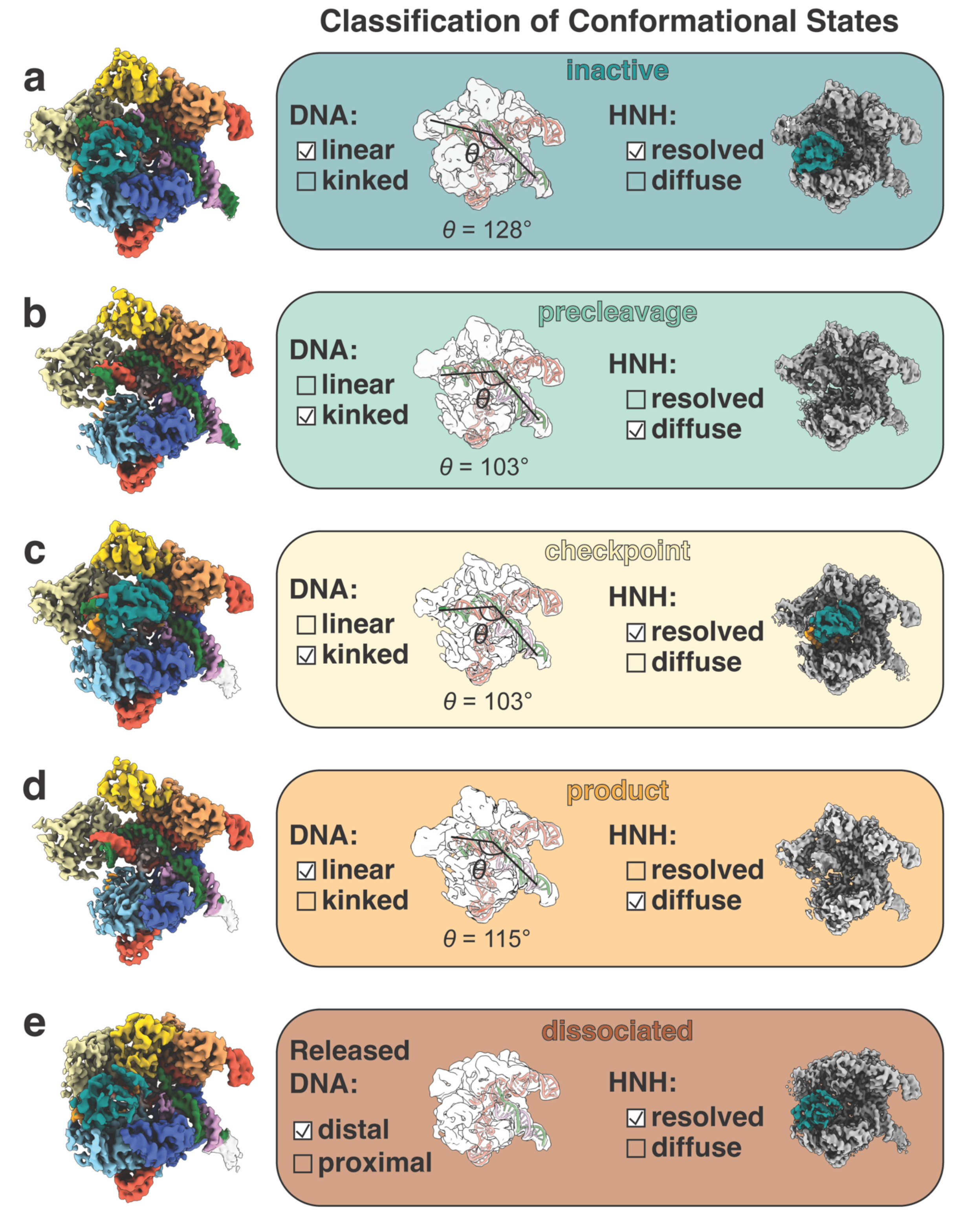


**Extended Data Figure 4. Conformational state assignment criteria. (a-e)** Metrics used to manually inspect and classify subclasses into the inactive, pre-cleavage, checkpoint, product and dissociated states. **(a)** In the inactive state, the DNA duplex adopts a linear conformation and the HNH domain is clearly resolved and remains docked onto the NUC lobe. **(b)** In the pre-cleavage state, the DNA adopts a kinked conformation and triggers undocking of the HNH domain. HNH is flexible as it reorganizes and is not resolved. The TS remains intact. **(c)** In the checkpoint conformation, the DNA is still kinked but the TS has been cleaved. The HNH domain is clearly resolved and has undocked from the cut TS. **(d)** In the product state, the DNA has been cleaved and the PAM-distal duplex adopts an intermediate conformation in-between the linear and kinked states observed in the inactive and pre-cleavage states. HNH is flexible and is not resolved in this state. **(e)** In the dissociated state, the PAM-distal DNA is released and the HNH domain is clearly observed to be reset into the position as observed in the binary complex.

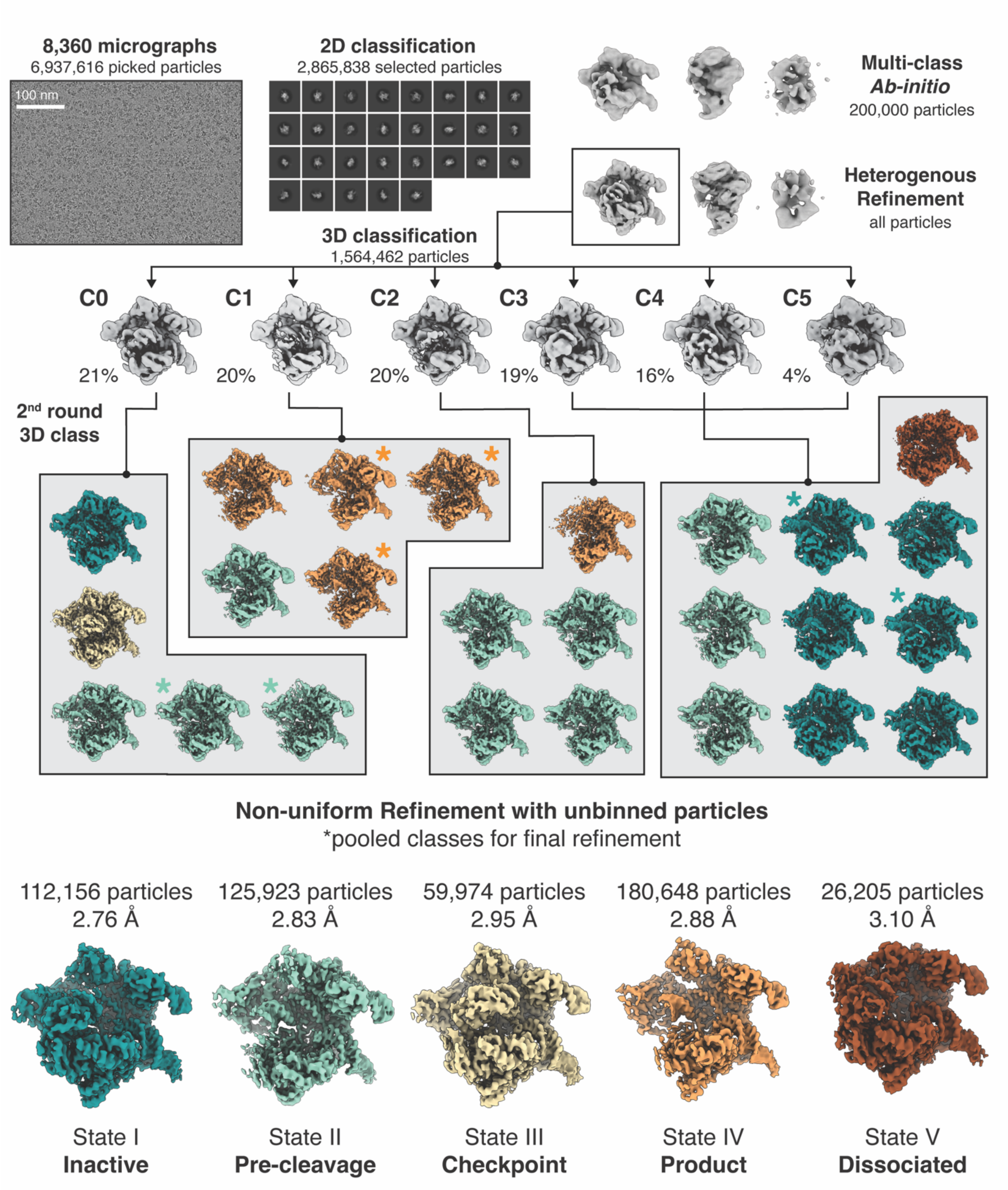


**Extended Data Figure 5. Cryo-EM data analysis workflow.**

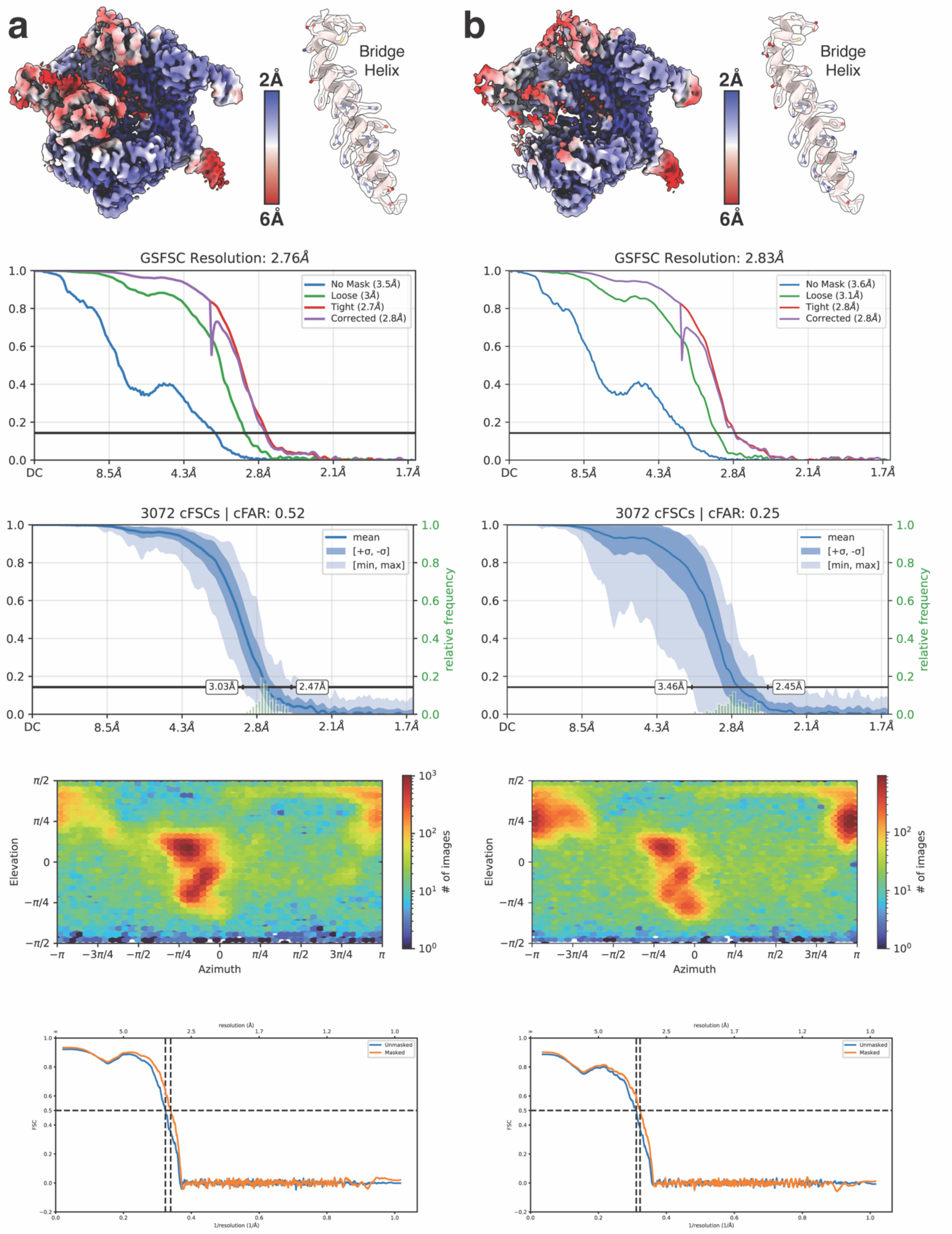

a
b
Bridge Helix
2Å
6Å
GSFSC Resolution: 2.76Å
No Mask (3.5Å)
Loose (3Å)
Tight (2.7Å)
Corrected (2.8Å)
GSFSC Resolution: 2.83Å
No Mask (3.6Å)
Loose (3.1Å)
Tight (2.8Å)
Corrected (2.8Å)
DC
8.5Å
4.3Å
2.8Å
2.1Å
1.7Å
3072 cFSCs | cFAR: 0.52
3.03Å
2.47Å
3072 cFSCs | cFAR: 0.25
3.46Å
2.45Å
mean
[+σ, -σ]
[min, max]
relative frequency
Elevation
Azimuth
### of images
Unmasked
Masked
resolution (Å)
1/resolution (1/Å)
FSC

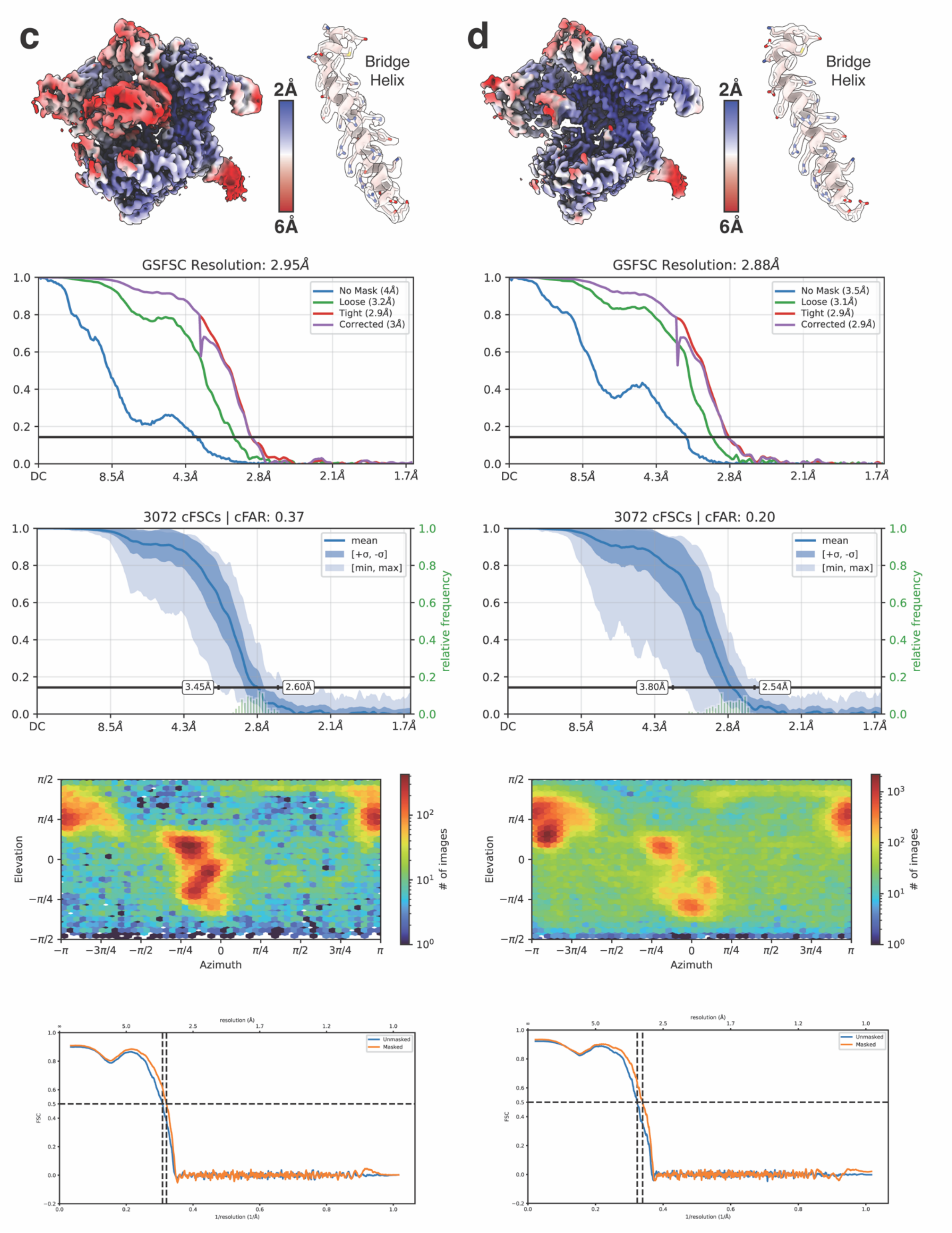

c
d
Bridge Helix
2Å
6Å
GSFSC Resolution: 2.95Å
GSFSC Resolution: 2.88Å
No Mask (4Å)
Loose (3.2Å)
Tight (2.9Å)
Corrected (3Å)
No Mask (3.5Å)
Loose (3.1Å)
Tight (2.9Å)
Corrected (2.9Å)
DC
8.5Å
4.3Å
2.8Å
2.1Å
1.7Å
3072 cFSCs | cFAR: 0.37
3072 cFSCs | cFAR: 0.20
mean
[+σ, -σ]
[min, max]
relative frequency
3.45Å
2.60Å
3.80Å
2.54Å
Elevation
Azimuth
### of images
resolution (Å)
Unmasked
Masked
FSC
1/resolution (1/Å)

**e**

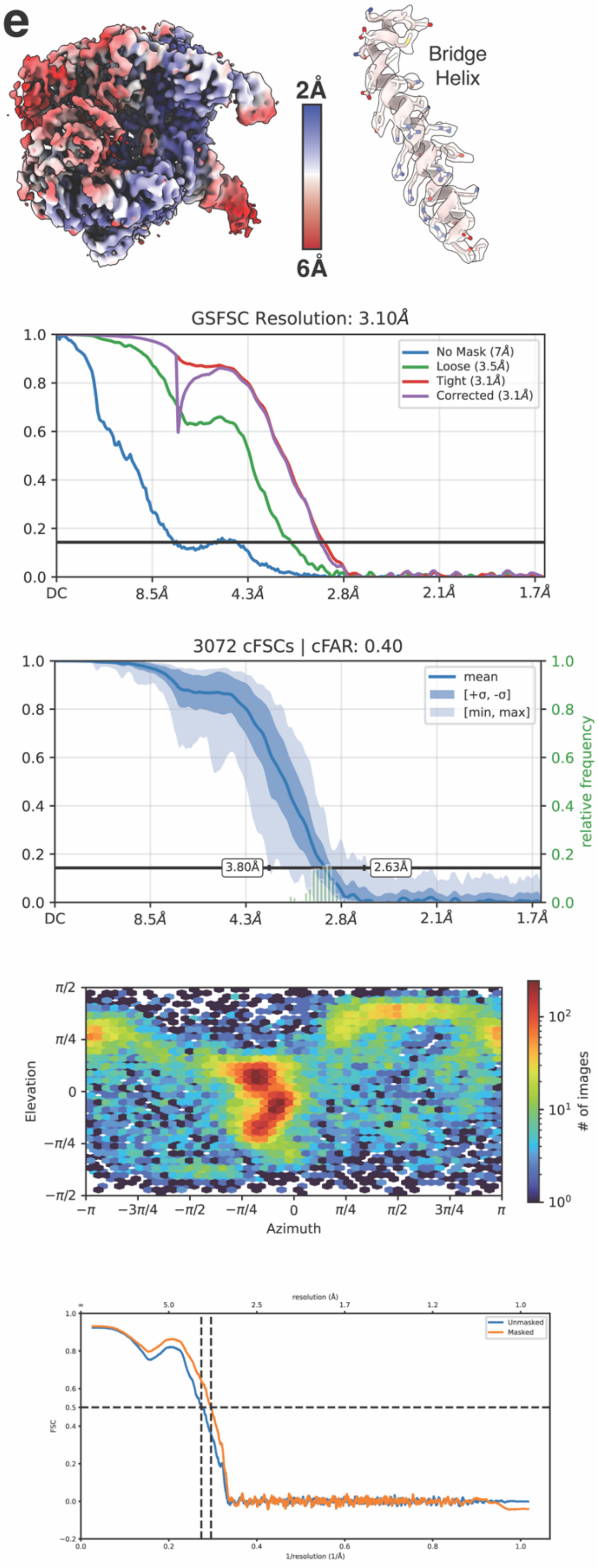

2Å
6Å
Bridge Helix
GSFSC Resolution: 3.10Å
No Mask (7Å)
Loose (3.5Å)
Tight (3.1Å)
Corrected (3.1Å)
DC
8.5Å
4.3Å
2.8Å
2.1Å
1.7Å
3072 cFSCs | cFAR: 0.40
mean
[+σ, -σ]
[min, max]
relative frequency
3.80Å
2.63Å
Elevation
Azimuth
### of images
resolution (Å)
Unmasked
Masked
FSC
1/resolution (1/Å)

**Extended Data Figure 6. Map and model quality assessment. (a-e)** From top to bottom: local resolution of cryo-EM reconstructions, representative map quality showing resolution of bridge helix sidechains, FSC curves, circular FSC curves, angular distribution plots, and map-to-model FSC curves for **(a)** the inactive state, **(b)** the pre-cleavage state, **(c)** the checkpoint state, **(d)** the product state, and **(e)** the dissociated state.

| Data Collection | State I | State II | State III | State IV | State V |
|---|---|---|---|---|---|
| Magnification | 81,000 | 81,000 | 81,000 | 81,000 | 81,000 |
| Voltage (kV) | 300 | 300 | 300 | 300 | 300 |
| Electron exposure ($e^-$/Å$^2$) | 80 | 80 | 80 | 80 | 80 |
| Defocus range (μM) | -1.2 to -2.2 | -1.2 to -2.2 | -1.2 to -2.2 | -1.2 to -2.2 | -1.2 to -2.2 |
| Pixel size (Å) | 0.8332 | 0.8332 | 0.8332 | 0.8332 | 0.8332 |
| Symmetry imposed | $C_1$ | $C_1$ | $C_1$ | $C_1$ | $C_1$ |
| Initial particle images | 1,564,462 | 1,564,462 | 1,564,462 | 1,564,462 | 1,564,462 |
| Final particle images | 112,156 | 125,923 | 59,974 | 180,648 | 26,205 |
| Map resolution (Å) | 2.76 | 2.83 | 2.95 | 2.88 | 3.10 |
| FSC threshold | 0.143 | 0.143 | 0.143 | 0.143 | 0.143 |
| Map resolution range (Å) | 2.5-7 | 2.5-7 | 2.5-7 | 2.5-7 | 2.5-7 |
| **Refinement** | **State I** | **State II** | **State III** | **State IV** | **State V** |
| Initial model (PDBID) | 6O0Z | 7S4V | 7Z4L | 7S4V | 4ZT0 |
| Model resolution (Å) | 2.9 | 3.1 | 3.1 | 3.2 | 3.4 |
| FSC threshold | 0.5 | 0.5 | 0.5 | 0.5 | 0.5 |
| Sharpening *B* factor (Å$^2$) | -61.3 | -65.0 | -57.1 | -65.1 | -46.6 |
| Model Composition | | | | | |
| Non-hydrogen atoms | 12,880 | 11,498 | 13,801 | 12,573 | 12,644 |
| Protein residues | 1,356 | 1,167 | 1,337 | 1,190 | 1,342 |
| Nucleotides | 149 | 143 | 135 | 135 | 122 |
| Ligands | 1 | 0 | 0 | 0 | 0 |
| Mean *B* factors (Å$^2$) | | | | | |
| Protein | 113.76 | 101.31 | 107.82 | 105.55 | 122.04 |
| Nucleotides | 129.21 | 109.23 | 95.62 | 108.29 | 124.34 |
| Ligands | 85.04 | N/A | N/A | N/A | N/A |
| R.m.s. deviations | | | | | |
| Bond lengths (Å) | 0.004 | 0.005 | 0.004 | 0.006 | 0.005 |
| Bond angles (°) | 0.800 | 0.874 | 0.840 | 1.178 | 0.895 |
| Validation | | | | | |
| MolProbity score | 1.50 | 1.50 | 1.42 | 1.65 | 1.52 |
| Clash score | 3.74 | 4.92 | 4.28 | 6.85 | 4.92 |
| Poor rotamers (%) | 1.09 | 0.94 | 0.58 | 0.66 | 0.43 |
| Ramachandran plot | | | | | |
| Favored (%) | 95.56 | 96.34 | 96.69 | 96.02 | 96.09 |
| Allowed (%) | 4.44 | 3.66 | 3.31 | 3.98 | 3.91 |
| Disallowed (%) | 0 | 0 | 0 | 0 | 0 |

**Extended Data Table 1. Cryo-EM data collection and refinement statistics.**

| Name | Sequence (5'-3') | Source |
|---|---|---|
| 55bp_TS | AGCTGACGTTTGTACTCCAGCGTCTCATCTTTATGCGTCAGCAGAGATTTCTGCT | IDT |
| 55bp_NTS | AGCAGAAATCTCTGCTGACGCATAAAGATGAGACGCTGGAGTACAAACGTCAGCT | IDT |
| 20nt_sgRNA | GGCGCAUAAAGAUGAGACGCGUUUUAGAGCUAGAAAUAGCAAGUUAAAAUAAGGCUAGUCCGUUAUCAACUUGAAAAAGUGGCACCGAGUCGGUGCUUUU | Genscript |
| 17nt_sgRNA | GCAUAAAGAUGAGACGCGUUUUAGAGCUAGAAAUAGCAAGUUAAAAUAAGGCUAGUCCGUUAUCAACUUGAAAAAGUGGCACCGAGUCGGUGCUUUU | Genscript |
| 16nt_sgRNA | CAUAAAGAUGAGACGCGUUUUAGAGCUAGAAAUAGCAAGUUAAAAUAAGGCUAGUCCGUUAUCAACUUGAAAAAGUGGCACCGAGUCGGUGCUUUU | Genscript |
| 15nt_sgRNA | AUAAAGAUGAGACGCGUUUUAGAGCUAGAAAUAGCAAGUUAAAAUAAGGCUAGUCCGUUAUCAACUUGAAAAAGUGGCACCGAGUCGGUGCUUUU | Genscript |
| 14nt_sgRNA | UAAAGAUGAGACGCGUUUUAGAGCUAGAAAUAGCAAGUUAAAAUAAGGCUAGUCCGUUAUCAACUUGAAAAAGUGGCACCGAGUCGGUGCUUUU | Genscript |
| 15mm1_sgRNA | GAUAAAGAUGAGACGCGUUUUAGAGCUAGAAAUAGCAAGUUAAAAUAAGGCUAGUCCGUUAUCAACUUGAAAAAGUGGCACCGAGUCGGUGCUUUU | Genscript |
| 15mm2_sgRNA | CGAUAAAGAUGAGACGCGUUUUAGAGCUAGAAAUAGCAAGUUAAAAUAAGGCUAGUCCGUUAUCAACUUGAAAAAGUGGCACCGAGUCGGUGCUUUU | Genscript |

**Extended Data Table 2. DNA and sgRNA sequences used in this study.**
